# The SARS-CoV-2 replication-transcription complex is a priority target for broad-spectrum pan-coronavirus drugs

**DOI:** 10.1101/2021.03.23.436637

**Authors:** Setayesh Yazdani, Nicola De Maio, Yining Ding, Vijay Shahani, Nick Goldman, Matthieu Schapira

## Abstract

In the absence of effective treatment, COVID-19 is likely to remain a global disease burden.
Compounding this threat is the near certainty that novel coronaviruses with pandemic potential will emerge in years to come. Pan-coronavirus drugs – agents active against both SARS-CoV-2 and other coronaviruses – would address both threats. A strategy to develop such broad-spectrum inhibitors is to pharmacologically target binding sites on SARS-CoV-2 proteins that are highly conserved in other known coronaviruses, the assumption being that any selective pressure to keep a site conserved across past viruses will apply to future ones. Here, we systematically mapped druggable binding pockets on the experimental structure of fifteen SARS-CoV-2 proteins and analyzed their variation across twenty-seven α- and β-coronaviruses and across thousands of SARS-CoV-2 samples from COVID-19 patients. We find that the two most conserved druggable sites are a pocket overlapping the RNA binding site of the helicase nsp13, and the catalytic site of the RNA-dependent RNA polymerase nsp12, both components of the viral replication-transcription complex. We present the data on a public web portal (https://www.thesgc.org/SARSCoV2_pocketome/) where users can interactively navigate individual protein structures and view the genetic variability of drug binding pockets in 3D.

---

The coronavirus SARS-CoV-2 that emerged in late 2019 has so far caused over two million deaths worldwide. While the recent approval of the first vaccines is expected to terminate the pandemic nature of the epidemic, it is believed that COVID-19 will not be eradicated, emphasizing the need for drugs^1^. Available vaccines may be less effective against novel and future SARS-CoV-2 variants, a serious global health threat. Additionally, in years to come, novel coronavirus outbreaks are bound to emerge, as did SARS-CoV-1 in 2003, the MERS coronavirus in 2012 and SARS-CoV-2 in 2019. To simultaneously develop anti COVID-19 therapeutics and prepare for the next pandemic threat, SARS-CoV-2 drug discovery efforts should focus on the development of pan-coronavirus agents. A first strategy is to target host proteins that are essential for the replication of past coronaviruses and may therefore be essential for the replication of coronaviruses emerging in the future^2^. An alternative approach, and the focus of this work, is to identify drug binding sites in the SARS-CoV-2 proteome that are conserved across coronaviruses. An approved anti-COVID-19 drug binding such site could be rapidly deployed against future outbreaks of novel coronaviruses.

Recent reports provided valuable insight into the druggability of SARS-CoV-2 proteins^3^, on their variation in COVID-19 samples and on their divergence from SARS-CoV-1 and from a specific bat coronavirus^4^. The excellent accompanying COVID-3D web portal allows users to map mutations on SARS-CoV-2 protein structures. A powerful online interface for inspecting variations between the SARS-CoV-2 genome and other viruses was also developed^5^, but to our knowledge, no integrated analysis focusing on the genetic variability of drug binding pockets across the SARS-CoV-2 proteome was reported to date.

We analyzed the structures of all SARS-CoV-2 protein domains represented in the PDB. The proteins analyzed included non-structural proteins (nsp) nsp1, nsp3, nsp5, nsp9, nsp12, nsp13, nsp14 (a SARS-CoV-1 structure was used that shared 100% sequence identity with SARS-CoV-2 at the active site), nsp15, nsp16; structural proteins E (transmembrane domain), Spike; additional factors ORF3a, ORF7a, ORF8, ORF9b. We used the PocketFinder function^6^ in ICM (Molsoft, San Diego) to find potential druggable binding sites and assessed their druggability using SiteMap (Schrodinger, NY)^7^. The druggability evaluated with SiteMap was in general agreement with a previous work using a different method^3^ and varied widely from poorly druggable, such as the catalytic site of the endoribonuclease nsp15 (Druggability score [Dscore] < 0.8), to highly druggable, such as a functionally uncharacterized ectopic site of the methyltransferase nsp14 (Dscore >1.0) (Figure 1a).

**Figure 1:**
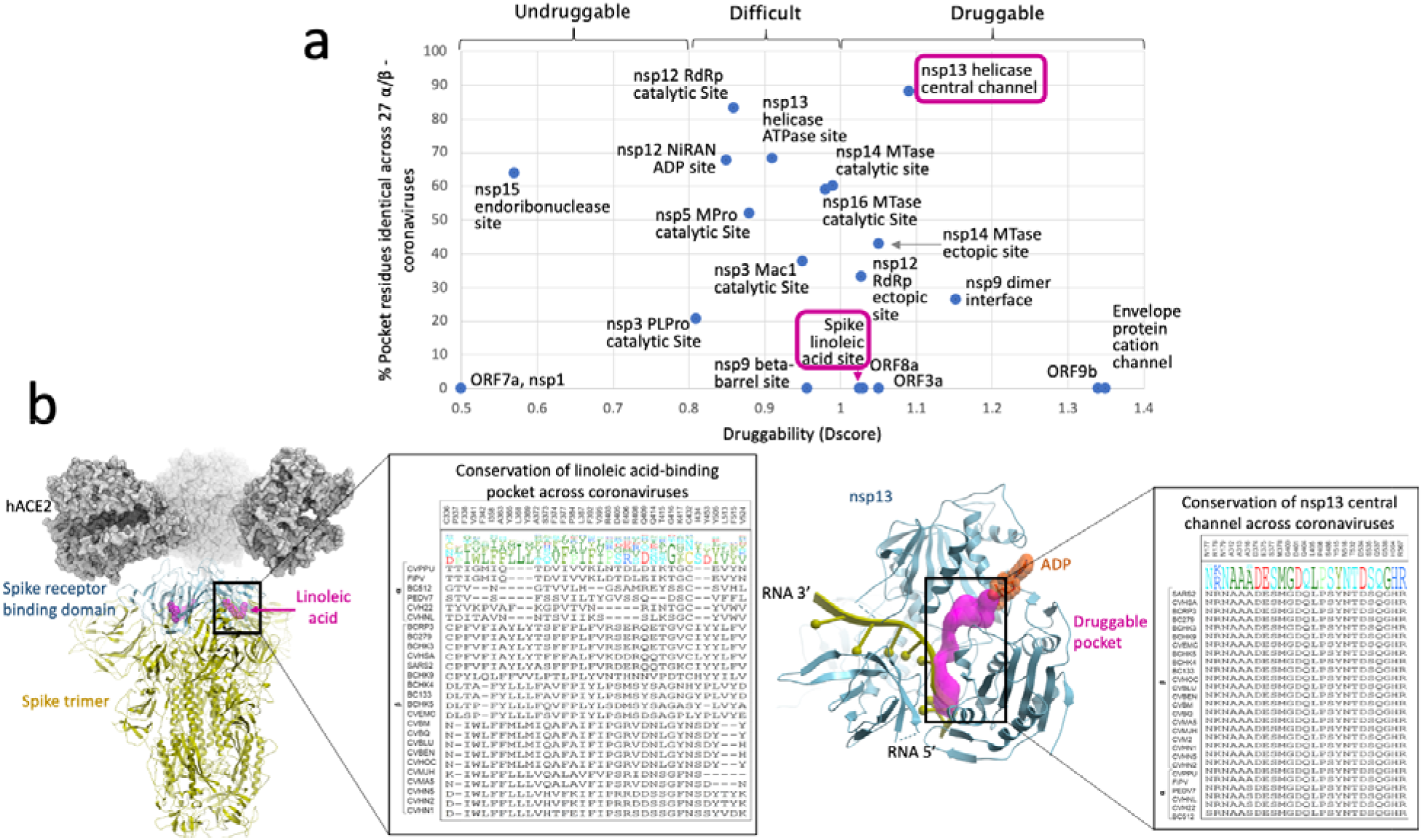
Druggability and conservation of drug binding sites in SARS-CoV-2 proteins. (a) For all binding sites, druggability scores calculated with SiteMap^7^ and conservation of pocket-lining residues across α- and β-coronaviruses are represented in a scatterplot. (b) Structural details and multiple sequence alignments are shown for the linoleic-acid binding site of the protein Spike (left, PDB code 6ZB4) which is poorly conserved, and a channel overlapping the RNA binding site of nsp13 (right, PDB code 6ZSL with RNA from structure 2XZL), which is highly conserved. Both sites are highlighted in the scatterplot (a). The full names of all viral genera are provided in Supplementary Table S1.

Since pathogenic coronaviruses responsible for mild or severe respiratory illness all belong to the genera alphacoronavirus (α-CoV) and betacoronavirus (β-CoV), we focused on these families to define the genetic variability of binding pockets found on the SARS-CoV-2 proteome^8^. Over 400 reviewed protein sequences of 58 α- and β-CoV were downloaded from UniProt (https://www.uniprot.org/). Automated sequence search retrieved homologs for most SARS-CoV-2 proteins, including all nsp proteins specified above, in 27 α- and β-CoV comprising zoonotic entries (bat and bovine CoVs), clinical and epidemic CoV isolates (SARS-CoV-1, MERS, HCoV-OC43, HCoV-HKU, HCoV-229E, HCoV-NL63). We performed multiple sequence alignments and determined the percent identity and conservation of amino-acids lining each binding pocket across all 27 coronaviruses where protein homologs were found (Figure 1a, Supplementary figure 1, Supplementary Table 1). We find that some binding sites are highly conserved while others are genetically unstable. For instance, binding of the essential fatty acid linoleic acid (LA) to the receptor binding domain of the SARS-CoV-2 Spike protein stabilizes the closed/locked state of Spike, inhibits Spike binding to ACE2 and synergizes with Remdesivir to reduce SARS-CoV-2 replication, suggesting that drugs targeting this site may be of therapeutic interest^9^. While the LA binding site is druggable (Dscore > 1.0), we find that sidechains lining LA are poorly conserved (0% conservation across 27 α- and β-coronaviruses), indicating that this site is not favorable for the discovery of pan-coronavirus inhibitors (Figure 1b). Conversely, a druggable channel partially overlapping with the RNA-binding site of the SARS-CcoV-2 helicase (nsp13) is highly conserved (>90% of residues lining the pocket are conserved across 27 α- and β-coronaviruses) (Figure 1b). Interestingly, nsp13 binds to the RNA-dependent RNA polymerase (nsp12) to functionally couple helicase and polymerase activities^10^ and we find that the catalytic site of nsp12 is also among the most conserved binding pockets in the SARS-CoV-2 proteome (>90% conservation across 27 α- and β-coronaviruses) (Figure 1a). These results support the notion of a strong selective pressure against genetic variability that may affect the function of the replication-transcription complex.

To further investigate genetic variability, we assessed mutation of residues lining our collection of binding pockets across thousands of SARS-CoV-2 samples from COVID-19 patients and found some correlation with amino-acid conservation across α- and β-coronavirus genera (supplementary Figure S2). For instance, the nsp13 druggable channel that is highly conserved across α- and β-coronaviruses is also conserved across SARS-CoV-2 samples: not a single residue lining this site was mutated across 15,000 samples. Conversely, the linoleic acid binding site of the Spike protein, which is poorly conserved across coronaviruses, is one of the most mutated across SARS-CoV-2 samples (38% residues mutated across 15,000 samples).

Our effort to systematically map the genetic variability of putative drug binding sites onto 3D structures of the SARS-CoV-2 proteome reveals medicinal chemistry strategies for the development of broad-spectrum coronavirus inhibitors. In particular, we find that pharmacologically targeting the viral replication-transcription complex is a promising avenue for the discovery of pan-coronavirus drugs. In this regard, compounds such as EIDD-280, an oral antiviral targeting the catalytic site of nsp12 and currently in phase II-III clinical trials against COVID-19^11^, represent promising drug candidates for current and future coronavirus pandemic threats. Similarly, targeting the RNA binding site of nsp13 is a priority – though underexplored – avenue that may benefit from recent advances on the chemical inhibition of RNA-binding proteins^12^.

## ASSOCIATED CONTENT

### Supporting Information

The following figures and tables are provided as supporting online information. Percent sequence identity and druggability of drug binding sites in the SARS-CoV-2 proteome represented in the PDB (Figure S1); mutation level of residues lining drug binding sites found in SARS-CoV-2 proteins in the PDB across >15,000 samples from COVID-19 patients and across 27 α- and β-coronavirus genera (Figure S2); conservation matrix of SARS-CoV-2 proteome represented in the PDB across 27 α- and β-coronaviruses (Table S1). Detailed methods are also provided.

## AUTHOR INFORMATION

### Author Contributions

The manuscript was written through contributions of all authors. All authors have given approval to the final version of the manuscript.

## ACKNOWLEDGMENT

We are grateful to Aled Edwards, Andrew Leach and Geoff Barton for insightful discussions at the outset of this work and to Dasha Redka for comments throughout the analysis. We are very grateful to GISAID and all the groups who shared their sequencing data, as detailed in https://github.com/roblanf/sarscov2phylo/tree/master/acknowledgements.

## ABBREVIATIONS

CoV: coronavirus
nsp: non-structural protein
PDB: Protein Data Bank
Dscore: Druggability Score

## TOC graphic

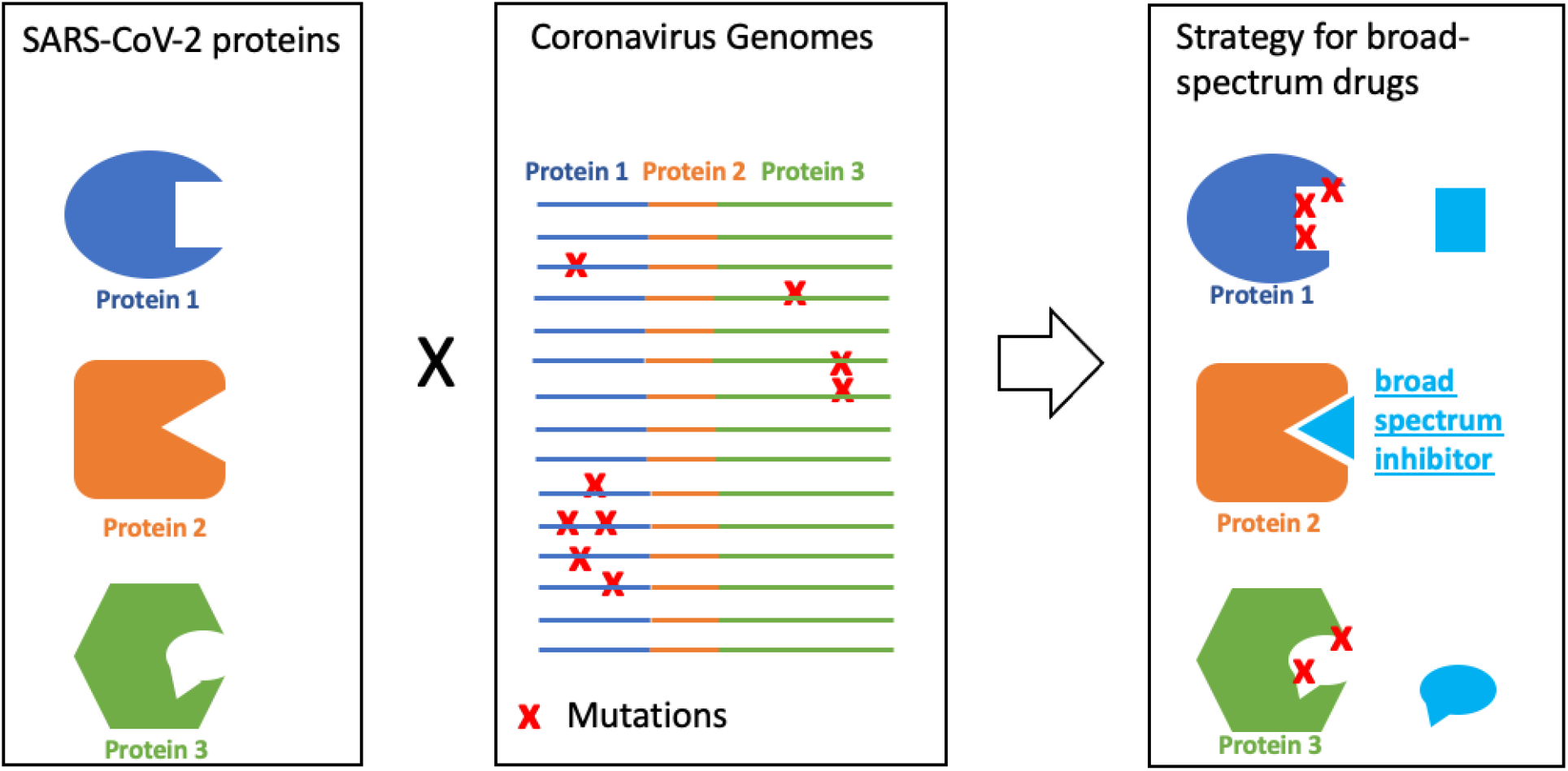

