## Supplementary information for "The SARS-CoV-2 replication-transcription complex is a priority target for broad-spectrum pan-coronavirus drugs"

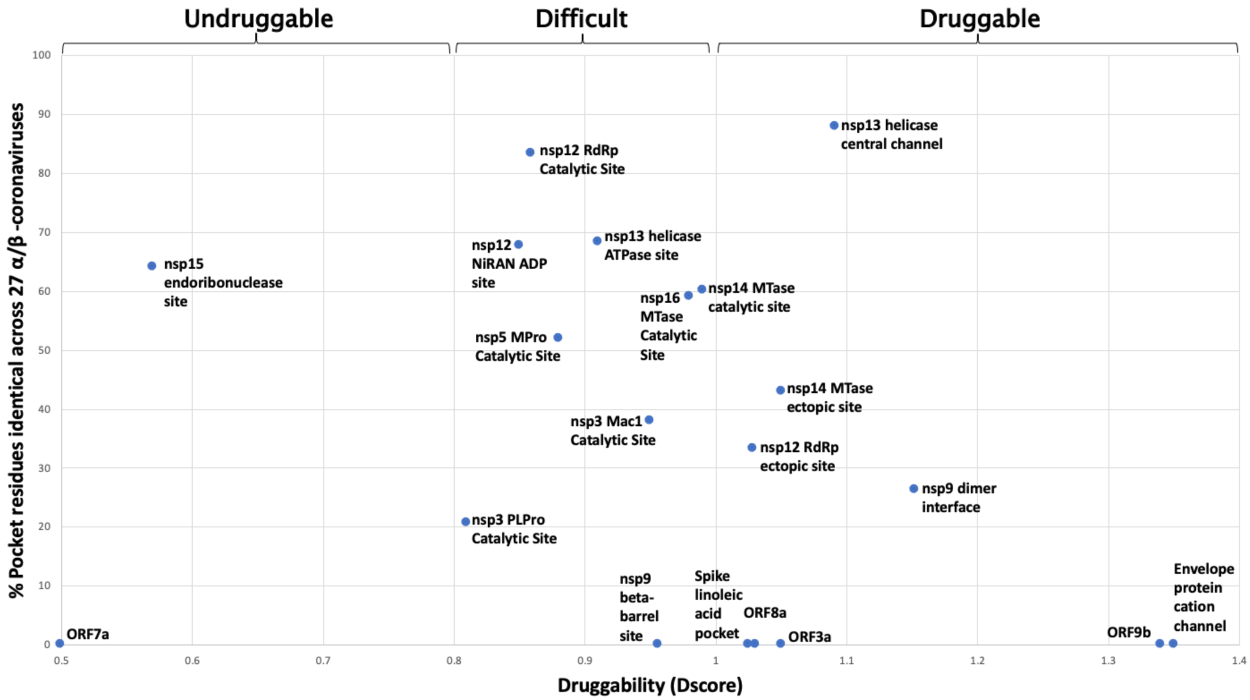


**Figure S1**: percent sequence identity and druggability of drug binding sites in the SARS-CoV-2 proteome represented in the PDB

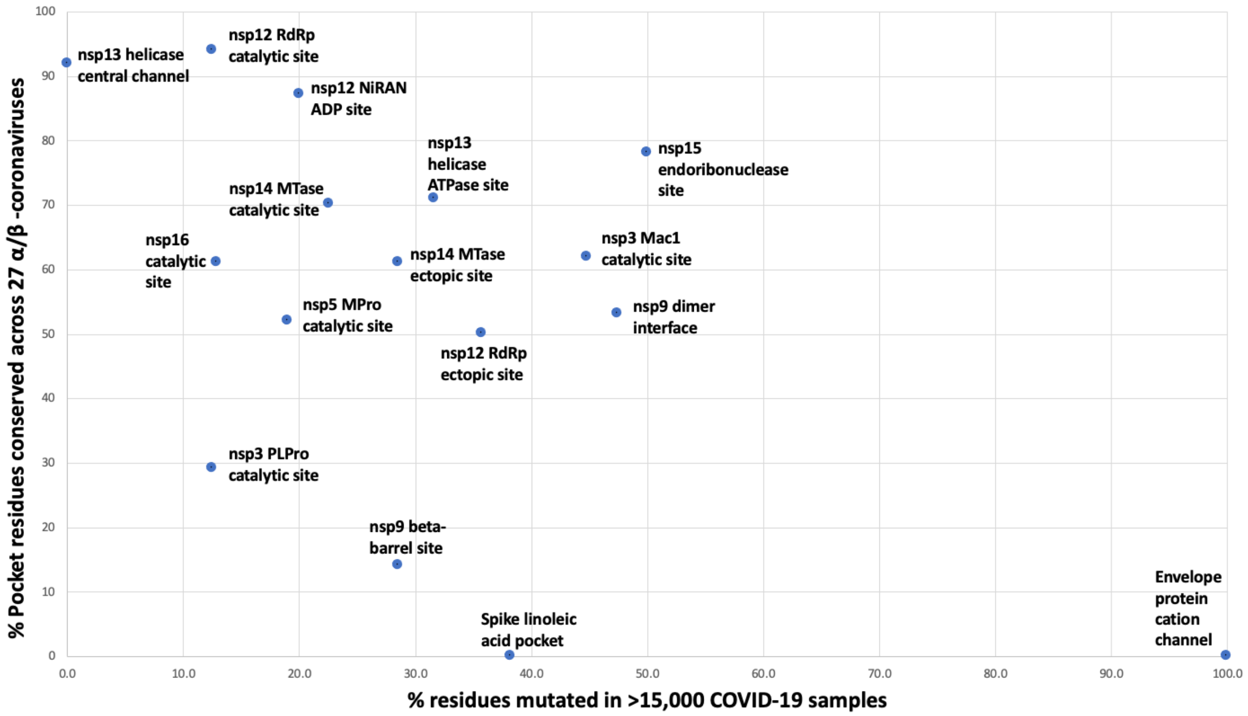


**Figure S2**: Mutation level of residues lining drug binding sites found in SARS-CoV-2 proteins in the PDB across >15,000 samples from COVID-19 patients and across 27 α- and β-coronavirus genera.

| | | | % sequence identity with SARS-CoV-2 at SARS-CoV-2 binding pockets | | | | | | | | | | | | | | | | | | | |
|---|---|---|---|---|---|---|---|---|---|---|---|---|---|---|---|---|---|---|---|---|---|---|
| | | | Non-structural Proteins | | | | | | | | | | | | | | Structural Proteins | | Accessory Proteins | | | |
| Genus | Organism | Entry | nsp3 PLPro Catalytic site | nsp3 Mac1 Catalytic site | nsp5 MPro Catalytic Site | nsp9 dimer interface | nsp9 beta-barrel site | nsp12 RdRp ectopic site | nsp12 RdRp catalytic Site | nsp12 NiRAN ADP site | nsp13 helicase ATPase site | nsp13 helicase central channel | nsp14 MTase catalytic site | nsp14 MTase ectopic site | nsp15 endoribonuclease site | nsp16 MTase Catalytic Site | Spike linoleic acid pocket | Envelope protein cation channel | ORF3a | ORF7a | ORF8 | ORF9b |
| β | **Human SARS coronavirus (SARS-CoV) (Severe acute respiratory syndrome coronavirus** | CVHSA | 100 | 97 | 100 | 100 | 100 | 100 | **100** | 100 | 100 | **100** | 100 | 100 | 100 | 100 | 79 | 100 | 80 | N/A | 17 | 100 |
| β | Bat coronavirus Rp3/2004 (BtCoV/Rp3/2004) (SARS-like coronavirus Rp3) | BCRP3 | 100 | 93 | 100 | 100 | 100 | 100 | **100** | 100 | 100 | **100** | 100 | 100 | 100 | 97 | 88 | 100 | 65 | N/A | 55 | 100 |
| β | Bat coronavirus HKU3 (BtCoV) (SARS-like coronavirus HKU3) | BCHK3 | 100 | 97 | 100 | 100 | 100 | 100 | **100** | 100 | 100 | **100** | 98 | 100 | 100 | 100 | 88 | 100 | 60 | N/A | 55 | 89 |
| β | Bat coronavirus 279/2005 (BtCoV) (BtCoV/279/2005) | BC279 | 96 | 97 | 100 | 100 | 100 | 100 | **100** | 100 | 100 | **100** | 100 | 100 | 100 | 100 | 88 | 75 | 65 | N/A | 44 | 100 |
| β | **Severe acute respiratory syndrome coronavirus 2 (2019-nCoV) (SARS-CoV-2)** | SARS2 | 100 | 100 | 100 | 100 | 100 | 100 | **100** | 100 | 100 | **100** | 100 | 100 | 100 | 100 | 100 | 100 | 100 | N/A | 100 | 100 |
| β | Bat coronavirus HKU9 (BtCoV) (BtCoV/HKU9) | BCHK9 | 42 | 86 | 76 | 58 | 43 | 79 | **98** | 93 | 87 | **100** | 78 | 79 | 86 | 82 | 26 | N/A | N/A | N/A | N/A | N/A |
| β | Bat coronavirus 133/2005 (BtCoV) (BtCoV/133/2005) | BC133 | 33 | 72 | 76 | 53 | 29 | 75 | **100** | 90 | 84 | **100** | 90 | 82 | 86 | 90 | 18 | N/A | N/A | N/A | N/A | N/A |
| β | Bat coronavirus HKU4 (BtCoV) (BtCoV/HKU4/2004) | BCHK4 | 33 | 72 | 76 | 53 | 29 | 75 | **100** | 90 | 84 | **100** | 90 | 79 | 86 | 90 | 18 | N/A | N/A | N/A | N/A | N/A |
| β | Bat coronavirus HKU5 (BtCoV) (BtCoV/HKU5/2004) | BCHK5 | 42 | 72 | 76 | 53 | 7 | 75 | **100** | 97 | 84 | **100** | 90 | 82 | 86 | 92 | 24 | N/A | N/A | N/A | N/A | N/A |
| β | **Middle East respiratory syndrome-related coronavirus (Human coronavirus EMC)** | CVEMC | 46 | 76 | 76 | 53 | 14 | 68 | **100** | 97 | 87 | **100** | 90 | 86 | 86 | 92 | 18 | N/A | N/A | N/A | N/A | N/A |
| β | Human coronavirus OC43 (HCoV-OC43) | CVHOC | 42 | 66 | 76 | 47 | 14 | 71 | **98** | 83 | 89 | **96** | 83 | 89 | 79 | 79 | 15 | N/A | N/A | N/A | N/A | N/A |
| β | Bovine coronavirus (strain Quebec) (BCoV) (BCV) | CVBQ | 42 | 66 | 76 | 47 | 14 | 71 | **98** | 80 | 89 | **96** | 83 | 89 | 79 | 69 | 15 | N/A | N/A | N/A | N/A | N/A |
| β | Bovine coronavirus (strain Mebus) (BCoV) (BCV) | CVBM | 42 | 66 | 76 | 47 | 14 | 71 | **98** | 80 | 89 | **96** | 83 | 89 | 71 | 79 | 15 | N/A | N/A | N/A | N/A | N/A |
| β | Bovine coronavirus (strain 98TXSF-110-LUN) (BCoV-LUN) (BCV) | CVBLU | 42 | 66 | 76 | 47 | 14 | 71 | **98** | 80 | 89 | **96** | 83 | 89 | 79 | 79 | 15 | N/A | N/A | N/A | N/A | N/A |
| β | Bovine coronavirus (strain 98TXSF-110-ENT) (BCoV-ENT) (BCV) | CVBEN | 42 | 66 | 76 | 47 | 14 | 71 | **98** | 80 | 89 | **96** | 83 | 89 | 79 | 79 | 15 | N/A | N/A | N/A | N/A | N/A |
| β | Human coronavirus HKU1 (isolate N2) (HCoV-HKU1) | CVHN2 | 46 | 66 | 76 | 47 | 21 | 71 | **98** | 87 | 87 | **96** | 85 | 79 | 79 | 82 | 15 | N/A | N/A | N/A | N/A | N/A |
| β | Human coronavirus HKU1 (isolate N5) (HCoV-HKU1) | CVHN5 | 46 | 66 | 76 | 47 | 21 | 71 | **98** | 87 | 87 | **96** | 83 | 79 | 79 | 82 | 15 | N/A | N/A | N/A | N/A | N/A |
| β | Human coronavirus HKU1 (isolate N1) (HCoV-HKU1) | CVHN1 | 46 | 66 | 76 | 47 | 21 | 71 | **98** | 87 | 87 | **96** | 83 | 79 | 79 | 82 | 15 | N/A | N/A | N/A | N/A | N/A |
| β | Murine coronavirus (strain 2) (MHV-2) (Murine hepatitis virus) | CVM2 | 46 | 62 | 76 | 53 | 14 | 71 | **98** | 87 | 89 | **96** | 78 | 86 | 79 | 77 | N/A | N/A | N/A | N/A | N/A | N/A |
| β | Murine coronavirus (strain JHM) (MHV-JHM) (Murine hepatitis virus) | CVMJH | 46 | 62 | 76 | 53 | 14 | 68 | **98** | 87 | 87 | **96** | 78 | 89 | 71 | 77 | 15 | N/A | N/A | N/A | N/A | N/A |
| β | Murine coronavirus (strain A59) (MHV-A59) (Murine hepatitis virus) | CVMA5 | 46 | 62 | 76 | 53 | 14 | 71 | **98** | 87 | 89 | **96** | 78 | 86 | 79 | 77 | 12 | N/A | N/A | N/A | N/A | N/A |
| α | Bat coronavirus 512/2005 (BtCoV) (BtCoV/512/2005) | BC512 | 29 | 66 | 57 | 42 | 7 | 68 | **90** | 83 | 74 | **92** | 80 | 61 | 79 | 79 | 0 | N/A | N/A | N/A | N/A | N/A |
| α | Porcine transmissible gastroenteritis coronavirus (strain Purdue) (TGEV) | CVPPU | 46 | 66 | 57 | 47 | 14 | 61 | **92** | 87 | 76 | **100** | 83 | 61 | 79 | 72 | 3 | N/A | N/A | N/A | N/A | N/A |
| α | Feline coronavirus (strain FIPV WSU-79/1146) (FCoV) | FIPV | 46 | 69 | 57 | 47 | 14 | 61 | **92** | 87 | 76 | **100** | 83 | 61 | 79 | 72 | 3 | N/A | N/A | N/A | N/A | N/A |
| α | Porcine epidemic diarrhea virus (strain CV777) (PEDV) | PEDV7 | 29 | 66 | 57 | 53 | 7 | 64 | **90** | 90 | 76 | **96** | 83 | 71 | 79 | 74 | 9 | N/A | N/A | N/A | N/A | N/A |
| α | Human coronavirus NL63 (HCoV-NL63) | CVHNL | 33 | 79 | 62 | 47 | 7 | 61 | **88** | 90 | 74 | **96** | 85 | 68 | 79 | 77 | 3 | N/A | N/A | N/A | N/A | N/A |
| α | Human coronavirus 229E (HCoV-229E) | CVH22 | 38 | 62 | 62 | 47 | 7 | 61 | **88** | 83 | 74 | **96** | 83 | 71 | 79 | 77 | 6 | N/A | N/A | N/A | N/A | N/A |

**Table S1:** Conservation matrix of SARS-CoV-2 proteome represented in the PDB across 27 α- and β- coronaviruses. SARS-CoV-2, SARS and MERS are highlighted in bold.

### METHODS:

#### Binding pocket detection:

Protein structures from the PDB were loaded in ICM (Molsoft, San Diego). Proteins were protonated, missing side-chains were built using a biased-probability Monte Carlo energy minimization simulation in the internal coordinates space, optimal positions of added polar hydrogens were generated, correct orientation of side-chain amide groups for glutamine and asparagine and most favourable histidine isomers were identified. The PocketFinder algorithm implemented in ICM, which uses a transformation of the Lennard-Jones potential to identify ligand binding envelopes regardless of the presence of bound ligands, was then applied (An et al. 2004, 2005). All PDB codes are provided in the accompanying web portal at https://www.thesgc.org/SARSCoV2_pocketome/

#### Druggablity score:

Protein structures were loaded in Maestro (Schrodinger, New York), and prepared using the default protein preparation wizard, which includes adjustment of protonation state and polar hydrogen rotameric state. Druggability scores (Dscores) were calculated with Schrodinger's SiteMap, where druggability of a binding pocket is calculated as a weighted function of volume, hydrophobicity and enclosure. Benchmark analysis demonstrated that binding pockets where extended experimental effort failed to identify drug-like ligands had a Dscore lower than 0.8 while experimentally druggable pockets had a Dscore higher than 1.0. Dscores between these values generally corresponded to challenging binding sites that could potentially be targeted by covalent inhibitors or by polar molecules that necessitated a pro-drug strategy (Halgren, 2009).

**Genetic variability of binding pockets across coronaviruses**:

Automated sequence search based on a full gapped optimal sequence alignment (Abagyan and Batalov, 1997) retrieved coronavirus homologs for most SARS-CoV-2 proteins. A multiple sequence alignment was generated using hierarchical clustering of the sequences based on sequence similarity calculated with the ZEGA alignment (a modification of the Needleman and Wunsch algorithm permitting zero gap-end penalties, ZEGA alignment) and Gonnet residue substitution matrix [gon92] (Gonnet et al. 1992, Abagyan and Batalov 1997). Residues with side-chain atoms within 2.8Å of the ligand binding envelope detected in ICM were extracted from the alignment and used to calculated % conservation and % identity.

**Genetic variability of binding pockets across SARS-CoV-2 samples:**

Over 15000 sequences marked as 'complete' and 'high coverage' submitted up to 31/7/20 were downloaded from GISAID. These sequences were then aligned to the reference genome (NC_045512.2 accession from NCBI), and the alignment was used to infer a maximum likelihood phylogenetic tree and a mutation history using parsimony (details of alignment, alignment filtering, tree inference, and mutation history inference can be found in (Turakhia, Thornlow, et al., 2020)). Alignment sites containing putative systematic sequencing errors were masked (details in (De Maio et al.; Turakhia, De Maio, et al., 2020)).
